# Cellular and synaptic adaptations of neural circuits processing skylight polarization in the fly

**DOI:** 10.1101/838300

**Authors:** Gizem Sancer, Emil Kind, Juliane Uhlhorn, Julia Volkmann, Johannes Hammacher, Tuyen Pham, Haritz Plazaola-Sasieta, Mathias F. Wernet

## Abstract

Specialized ommatidia harboring polarization-sensitive photoreceptors exist in the ‘dorsal rim area’ (DRA) of virtually all insects. Although downstream elements have been described both anatomically and physiologically throughout the optic lobes and the central brain of different species, little is known about their cellular and synaptic adaptations and how these shape their functional role in polarization vision. We have previously shown that in the DRA of *Drosophila melanogaster*, two distinct types of modality-specific ‘distal medulla’ cell types (Dm-DRA1 and Dm-DRA2) are post-synaptic to long visual fiber photoreceptors R7 and R8, respectively. Here we describe additional neuronal elements in the medulla neuropil that manifest modality-specific differences in the DRA region, including DRA-specific neuronal morphology, as well as differences in the structure of pre- or post-synaptic membranes. Furthermore, we show that certain cell types (medulla tangential cells and octopaminergic neuromodulatory cells) specifically avoid contacts with polarization-sensitive photoreceptors. Finally, while certain transmedullary cells are specifically absent from DRA medulla columns, other subtypes show specific wiring differences while still connecting the DRA to the lobula complex, as previously been described in larger insects. This hints towards a complex circuit architecture with more than one pathway connecting polarization-sensitive DRA photoreceptors with the central brain.

## Introduction

Retinotopic photoreceptor axon projections onto columnar microcircuits ensure an accurate representation of the visual world within the optic lobes of insects (Braitenberg 1967). In fruit flies, both light microscopic studies as well as 3D reconstruction of electron microscopic data revealed that these repetitive neural ensembles (called cartridges or columns, depending on the neuropil) contain a stereotypic set of both unicolumnar and multicolumnar cell types (Gao, Takemura et al. 2008, Morante and Desplan 2008, Takemura, Lu et al. 2008, Takemura, Bharioke et al. 2013, Nern, Pfeiffer et al. 2015, Takemura, Xu et al. 2015). Little is known about differences between these neighboring columnar structures (like cellular composition, synaptic strength, and distribution) (Takemura, Kinoshita et al. 2005, Takemura and Arikawa 2006, Jagadish, Barnea et al. 2014, Karuppudurai, Lin et al. 2014, Takemura, Xu et al. 2015). Such differences are of particular interest since most insect eyes analyzed to date are in fact retinal mosaics containing molecularly and morphologically distinct ommatidial subtypes (for review: (Wernet, Perry et al. 2015)). Stochastic expression of different Rhodopsins within long visual fiber photoreceptors creates a mosaic with two (flies, locusts) or three (bees, butterflies) randomly distributed subtypes that mediate colour vision (Wakakuwa, Kurasawa et al. 2005, Wernet, Mazzoni et al. 2006, Perry, Kinoshita et al. 2016). Additionally, virtually all insect species also manifest specialized ommatidia in the ‘dorsal rim area’ (DRA; for a list of abbreviations, see supplemental table 1) of the adult eye, harboring photoreceptors that all express the same Rhodopsin while being specialized for the detection of polarized skylight (for review: (Labhart and Meyer 1999)), a stimulus that is of great importance for many navigating insects (Heinze 2017) (Fig. 1a).

**Fig. 1:**
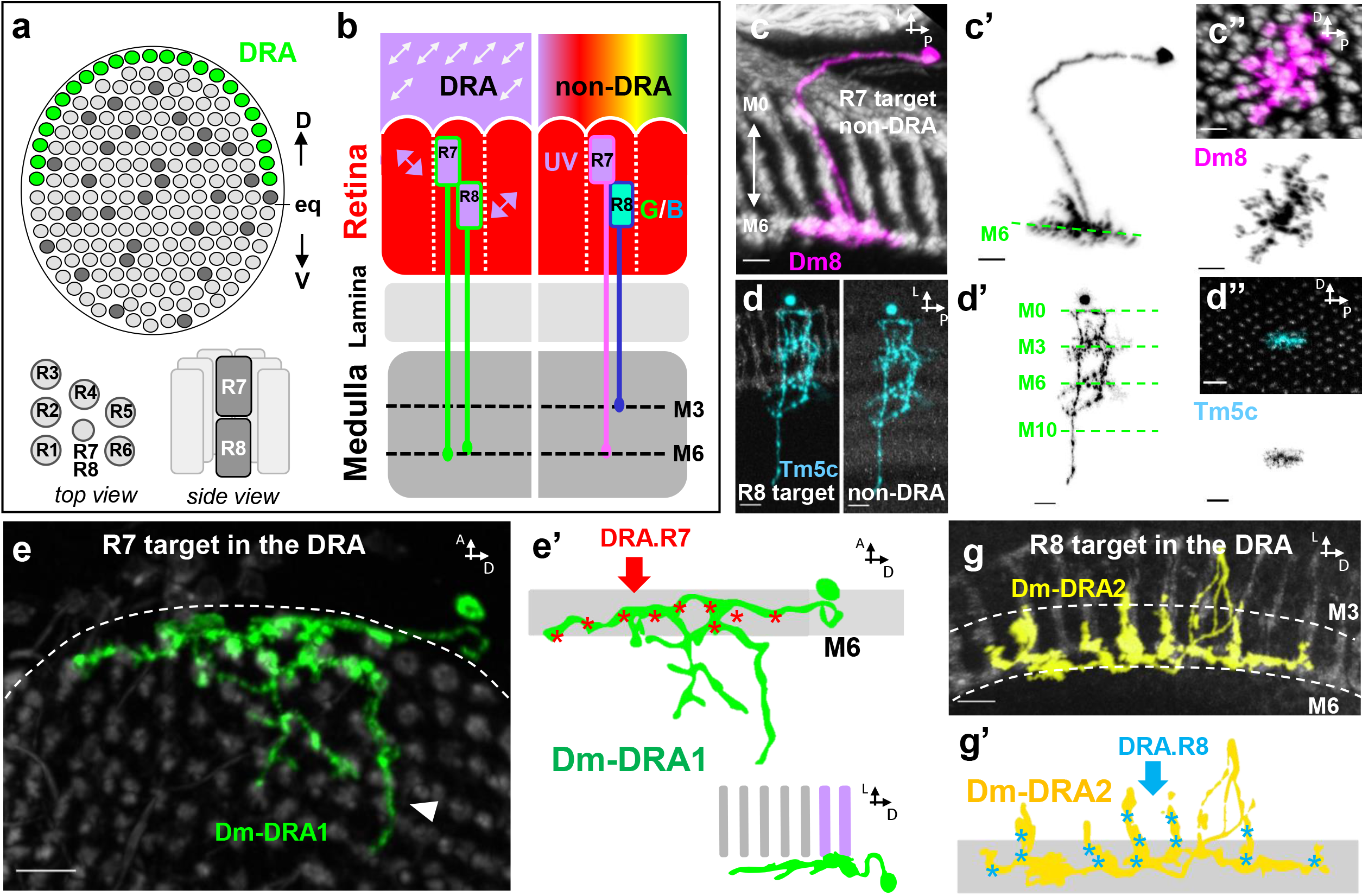
Modality-specific cell types in the DRA region of the fly visual system. **a** Top: Schematic representation of three ommatidial subtypes of the fly retina: stochastically distributed ‘pale’ and yellow’ (light and dark grey), as well as DRA ommatidia (green) in the ‘dorsal rim area’. D = dorsal; V = ventral; eq = equator. Bottom left: Top view onto a section through one ommatidium with outer photoreceptor (R1-6) rhabdomeres surrounding R7 and R8. Bottom right: Side view onto the same schematic ommatidium with R7 rhabdomeres located on top of R8. **b** Schematic description of the fly visual system. Individual ommatidial units within the retina (red) form repetitive units (separated by dashed lines) contain a full complement of eight photoreceptor neurons that project to the lamina neuropil (R1-6), or the medulla neuropil (R7, R8). Only in polarization-sensitive DRA ommatidia (left), both R7 and R8 terminate in deep medulla layer M6, whereas color-sensitive non-DRA R8 always terminate in the more distal layer M3 (right). **c** Single cell clone of the main synaptic target of non-DRA R7 cells (amacrinelike distal medulla cell type Dm8 in magenta) located in layer M6, seen both from the dorsal side (C’) as well as from a lateral view (c’’). **d** Main synaptic target of non-DRA R8 cells (transmedulla cell type Tm5c in cyan, bottom), with processes in several medulla layers. **e** Single cell clone of Dm-DRA1 (green), a modality-specific cell type in the DRA layer M6 with ‘deep projections’ (arrowhead) avoiding contacts with color-sensitive photoreceptor terminals. **f** Dm-DRA1 receives synaptic input from DRA.R7 cells specifically (red asterisks). **g** Single cell clone of Dm-DRA2 (yellow), another modality-specific cell type in the DRA layer M6. **h** Dm-DRA2 receives synaptic input from DRA.R8 cells specifically along vertical projections (light blue asterisks). Scale bars: 5μm in (c) and (g), 10 μm in (d) and (e).

The fruit fly *Drosophila melanogaster* serves as an attractive model system for investigating the structure and function of DRA circuitry, using the vast arsenal of molecular genetic tools available for this model system (Meinertzhagen and Lee 2012, Simpson and Looger 2018). Importantly, fly DRA ommatidia have been described in great detail (Wada 1971, Wada 1974, Wunderer and Smola 1982, Wernet, Labhart et al. 2003, Weir, Henze et al. 2016), behavioral responses to linearly polarized light have been demonstrated for flies (Hardie 1984, Vonphilipsborn and Labhart 1990, Weir and Dickinson 2012, Wernet, Velez et al. 2012, Velez, Gohl et al. 2014, Velez, Wernet et al. 2014, Warren, Weir et al. 2018), as well as physiological responses to linearly polarized light (Weir, Henze et al. 2016). Only in the DRA region of the fly visual system, both long visual fiber photoreceptors R7 and R8 both express the same UV Rhodopsin and acquire high polarization-sensitivity through the untwisted design of their light-gathering rhabdomeres. Importantly, rhabdomeres of R7 and R8 from the same ommatidium are oriented perpendicular to each other, making these cells maximally sensitive to orthogonal e-vector orientations, resulting in polarization-opponent analyzer pairs. Along the DRA, ommatdial analyzer directions change gradually, forming a fan-shaped array of polarization detectors (Weir, Henze et al. 2016). Only in DRA ommatidia, R7 and R8 axons terminate in the same deep layer of the medulla neuropil (M6) (Strausfeld and Wunderer 1985, Fischbach and Dittrich 1989, Chin, Lin et al. 2014), whereas colour-sensitive counterparts terminate in distinct layers M3 (R8 photoreceptors) and M6 (R7 photoreceptors) (Fischbach and Dittrich 1989) (Fig. 1b). The segregation of colour-sensitive R7 versus R8 presynapses in different layers is believed to facilitate the establishment of synapses with different post-synaptic targets: computation of UV signals is initiated in layer M6, where ‘distal medulla’ cell type Dm8 serves as the major post-synaptic partner of 10-16 neighboring R7 photoreceptors (Gao, Takemura et al. 2008) (Fig. 1c). In contrast, R8 photoreceptors are sensitive to longer wavelengths form most of their synaptic connections in layer M3, where the most prominent target is the transmedulla cell type Tm5c (Gao, Takemura et al. 2008, Karuppudurai, Lin et al. 2014, Kulkarni, Ertekin et al. 2016) (Fig. 1d).

Polarization-sensitive circuit elements have been described throughout the optic lobes as well as the central brain of several insect species (for review: (el Jundi, Pfeiffer et al. 2014, Homberg 2015). Ultimately, different e-vector orientations of linearly polarized skylight become represented in a map-like arrangement of columnar units of the central complex (Heinze and Homberg 2007, Sakura, Lambrinos et al. 2008, Heinze and Reppert 2011), where they are integrated with other cues like chromatic information (el Jundi, Warrant et al. 2015). An important relay station between the optic lobes and the central complex is an optic glomerulus called the ‘anterior optic tubercle’ (AOTU), the likely site where information from the circadian clock and skylight polarization are integrated (Pfeiffer and Homberg 2007). This is particularly important for navigating insects that have an interest in keeping a straight course while the position of the sun (and the polarization pattern around it) changes over the course of the day. Although direct connection from the medulla towards the AOTU have been demonstrated in various insects (Homberg, Hofer et al. 2003, Pfeiffer and Kinoshita 2012, Omoto, Keles et al. 2017), it is likely that polarized light information is processed via other pathways as well, both in the medulla and the lobula complex neuropils (El Jundi and Homberg 2010). To date, the functional significance of such parallel processing streams remains poorly understood. Interestingly, little is known about how photoreceptor signals are computed by units post-synaptic to DRA long visual fiber photoreceptors in the medulla neuropil (Strausfeld and Wunderer 1985, el Jundi, Pfeiffer et al. 2011).

We have recently shown that medulla columns in the DRA region of the fly visual system contain two types of modality-specific ‘distal medulla’ (Dm) cell types (termed Dm-DRA1 and Dm-DRA2) that are specifically connected to either DRA.R7 cells (Fig. 1e,f) or DRA.R8 cells (Fig. 1g,h), respectively (Sancer, Kind et al. 2019). Importantly, these cells exclusively contact polarization-sensitive photoreceptors while specifically avoiding colour-sensitive photoreceptors. Outside the DRA region, Dm8 cells within layer M6 collect visual information from ~14 neighboring R7 photoreceptors (Gao, Takemura et al. 2008, Karuppudurai, Lin et al. 2014, Ting, McQueen et al. 2014). These Dm8 cells never overlap with Dm-DRA1 and Dm-DRA2 cells, resulting in a strict modality-specific boundary between DRA and non-DRA territories. Since Dm-DRA1 and Dm-DRA2 cells (like Dm8 cells) heavily overlap with their own kind, these cells collect signals from neighboring DRA ommatidia with slightly divergent preferred e-vector tuning (Weir, Henze et al. 2016). The signals collected by each type should differ by 90 degrees, due to the fact that Dm-DRA1 and Dm-DRA2 cells located at the same position along the DRA receive input only from DRA.R7 or DRA.R8, respectively, whose preferred e-vector orientations differ by 90 degrees. It remains unknown how information is compared between Dm-DRA1 and Dm-DRA2 cells, nor do we know who the post-synaptic elements are.

Here we present a systematic morphological characterization of cell types located within the DRA region of the optic lobes of *Drosophila* (for list of cell types studied, see supplemental table 2). We show that several ‘distal medulla’ (Dm cell types (Fischbach and Dittrich 1989, Nern, Pfeiffer et al. 2015) show DRA-specific morphologies and/or DRA-specific distribution of fluorescently labeled pre-versus postsynaptic membranes (Dm2 and Dm9, but not Dm4). Similarly, we show that several multicolumnar cell types specifically avoid contacting DRA photoreceptors, or invading DRA columns altogether (Mt11-like cells and Tdc2-expressing octopaminergic, neuromodulatory cells). Of the ‘transmedullary’ (Tm) cell types, which connect the distal medulla with the lobula complex, Tm5c and Tm20 located in the DRA are postsynaptic to polarization-sensitive long visual fiber photoreceptors, potentially demonstrating modality-specific differences in their R7-versus R8 specificity. In contrast, other Tm cell types like Tm5a,b appear to be specifically absent in the DRA region. Taken together, these data points towards the computation of polarized light information involving a modality-specific medulla-to-lobula circuit from photoreceptors towards the central brain, in analogy to what has been shown in larger insect species.

## Materials and methods

### Fly Strains

The flies were maintained on standard molasses-corn food at 25°C 12h light/dark cycle incubator unless otherwise mentioned.

#### driver lines

longGMR-Gal4 (all photoreceptors; BDSC#8605), LongGMR-LexA (all photoreceptors), *rh3*-Gal4^−137^ (Sancer, Kind et al. 2019), rh3-LexA (gift from C. Desplan), DRA.R8-Gal4 (Sancer, Kind et al. 2019), DRA.R8-LexA, GMR24F06 (Dm8 & Dm-DRA1, BDSC#49087), ort^c2b^-Gal4 (Dm8, Dm-DRA1+2, (Karuppudurai, Lin et al. 2014)), ort^C1-3^-LexADBD, ort^C2B^-dVP16AD (Dm8, Dm-DRA1+2, (Karuppudurai, Lin et al. 2014)); GMR26H07-GAL4 (Dm2, BDSC#50328), GMR23G11-GAL4 (Dm4, BDSC#49043), GMR21A12-Gal4 (Dm9, gift from C. Desplan), ort^C1a^-Gal4DBD#3, ok371-dVP16AD(Tm5c, (Karuppudurai, Lin et al. 2014)), ort^C1a^-Gal4DBD#3, pVP16AD^24G^ Tm5a,b, (Karuppudurai, Lin et al. 2014)), GMR33H10-Gal4 (Tm20, BDSC#49762), GMR71C10-Gal4 (Mt11-like, BDSC#39576), Tdc2-Gal4 (BDSC#9313), TH-Gal4 (BDSC#8848).

#### Cell labeling

UAS-mCD8::GFP (BDSC#5137), MCFO-1 (Nern, Pfeiffer et al. 2015)

#### Active zone or postsynaptic localization

UAS-brp^D3^:mKate (gift from N. Özel), UAS-DRep2:GFP (gift from Stefan Sigrist).

#### Activity GRASP experiments

UAS-nSyb-spGFP^1-10^, lexAop-CD4-spGFP^11^, lexAop-nSyb-spGFP^1-10^ and UAS-CD4-spGFP^11^(Macpherson, Zaharieva et al. 2015).

### Immunohistochemistry and imaging

Adult brain dissection was performed in ice-cold S2 cell culture medium (Schneider’s Insect Medium, Sigma Aldrich, #S0146) and brains were fixed with 4% PFA (v/w) in PBS for 20 – 30 minutes at room temperature. After 3 times washing with PBS-T [PBS with 0.5% (v/v) Triton X-100 (Sigma Aldrich, # X-100)] fixed brains were incubated with primary antibody containing 10% Normal Donkey Serum in 0.4% PBS-T overnight at 4°C. Following three times washing with PBS-T, brains were incubated with secondary antibody solution containing 10% Normal Donkey Serum in 0.4% PBS-T overnight. After 3 times 15 minutes washing brains with PBS were mounted in Vectashield H-1000 (Vector Laboratory, Burlingame, CA) anti-fade mounting medium for confocal microscopy. The primary antibodies used in this study were anti-chaoptin (24B10) mouse (1:250), anti-BRP (nc82) mouse (1:50), anti-HA rat (1:250), anti-CD4 rabbit (1:600), anti-dsRed rabbit (1:500), anti-FLAG chicken (1:1000), anti-GFP mAb rat (1:500), anti-GFP pAb rabbit(1:1000), anti-GFP pAb goat (1:1000), anti-Ncad (DN-Ex #8) rat (1:100), V5 epitope tag antibody Dylight™ 549 conjugated rabbit (1:1000). Secondary antibodies were diluted at 1:500 and were as follows: anti-chicken Cy™5 donkey, anti-rabbit BV480 donkey, anti-goat Cy™5 donkey, anti-rat Cy™5 donkey, anti-rabbit Cy™5 donkey, Jackson Immuno Research anti-rabbit Cy™3 donkey, anti-mouse Alexa Fluor^®^ 594 donkey, anti-mouse Cy™5 donkey, anti-rabbit Alexa Fluor^®^ 488 donkey, anti-rat Alexa Fluor^®^ 488 donkey.

A Leica SP8 confocal microscope equipped with a white light laser and two HyD detectors was used. Image stacks were acquired in resolution of 1024×1024×0.5μm with a 63x lens.

### Single cell clones and MCFO

To obtain single cell clones and reveal the morphology and relative position of individual neurons in the adult visual system, hsFLP and MCFO-1 (Nern, Pfeiffer et al. 2015) were used. 3 days-old flies were incubated in vials in a 37°C water bath for 10-60 minutes 3 days prior to fly brain dissection to induce flippase (FLP). To allow the expression of the reporter, the flies were kept over 3 days at 25° C. Dissection and staining occurred as described above.

### Activity GRASP

Flies were grown in a 25° C, 12h-12h dark-light cycle incubator in normal vials and transferred to custom-made UV-transparent Plexiglas tubes [wall thickness: 4mm, From RELI KUNSTSTOFFE (Kunststoffhersteller in Erkner, Brandenburg) (Gewerbegebiet zum Wasserwerk 16, 15537 Erkner)] before light induction. 1-day old flies were kept in a 25°C, 20 h – 4 h light-dark cycle custom-made light box (including unpolarized UV light) for 3 days to ensure syfficient photoreceptor activation in the DRA.

Dissection & staining occurred as described above. Brains were stained with polyclonal GFP (anti-GFP goat pAB) and monoclonal GFP (anti-GFP rat mAB) antibody to visualize postsynaptic cells and GRASP signal, respectively. Postsynaptic cells were visualized by staining with CD4 antibody.

### Morphology and characteristics of the cells

Analysis and post-processing were done using IMARIS software and FIJI. Single cell clones from MCFO data stacks were obtained by using the Surface and masking function in IMARIS. Contact points for neurons were determined according to photoreceptor surface and cell surface contact points and reviewed with fluorescent signal

### Columnar layer analysis

A row of photoreceptors containing DRA and non-DRA columns was extracted by using the 3D Crop function in IMARIS and a snapshot was loaded in to FIJI. With the segmented line tool (line width 25) the induvial columns were traced from M0 to M10 (based on 24B10 and Ncad staining). The average intensity of the GFP channel along the segmented line was measured with the plot profile tool and the corresponding csv files were loaded into R. The segmented line length (which corresponds to the length of the column from M0-M10) was normalized. Graphs were plotted with R and the individual medulla layers were annotated based on their medulla coverage along the distal-proximal medulla axis described by Fischbach and Dittrich (Fischbach and Dittrich 1989).

## Results

### Morphology and connectivity of different ‘Distal medulla’ (Dm) cell types in the DRA

In order to characterize both the single-cell morphology as well as the synaptic connectivity of cell types in the DRA region of the medulla, we first screened a collection of previously published Gal4 drivers specifically expressed in different ‘distal medulla’ (Dm) cell types (Nern, Pfeiffer et al. 2015). Besides Dm-DRA1 and Dm-DRA2 (both labeled by different Dm8-specific driver lines (Sancer, Kind et al. 2019)), only Dm2 cells appeared to be modality-specific, i.e. any given single cell clone located at the dorsal edge of the medulla never contacted both DRA- and non-DRA photoreceptors (Fig. 2a, Supplemental Fig. S1). Since Dm2 cells usually cover only ~2 medulla columns in layer M6 (Nern, Pfeiffer et al. 2015), their morphology was rather uniform across the medulla (DRA and non-DRA). In the DRA, GFP signal from Dm2 cell processes formed a rather sharp peak in layer M6, whereas non-DRA Dm2 cell signal was flatter with a longer tail towards more distal layers (Fig. 2 a,b). In agreement with this, the overall intensity peak of DRep2:GFP signals from Dm2 cells (a marker for labeling putative post-synaptic membranes) was shifted towards layer M6 when compared to non-DRA Dm2 cells Fig. 2c). Interestingly, presynaptic signals (visualized using brp^D3^:mKate) in M6 were reduced in DRA columns, when compared to non-DRA signals (Fig. 2d). The activity-dependent version of GRASP (see materials & methods) revealed reconstituted GFP signals (and therefore potential synaptic connections) between photoreceptors (presynaptic) and Dm2 cells (postsynaptic), both in DRA and non-DRA columns (Fig. 2e). In non-DRA columns, these signals spread from M6 to ~M3, with a clear contribution from R7 cells (rh3 > Dm2 GRASP; Fig. 2f). The signal was also detectable into the DRA where it was more restricted to layer M6 (arrowhead in Fig 2f,g). An R8 contribution to the signals there (also labeled by rh3-Gal4) appeared unlikely due to a lack of signal using a DRA.R8-specific driver (Sancer, Kind et al. 2019) (DRA.R8 > Dm2 GRASP; Fig. 2g).

**Fig. 2:**
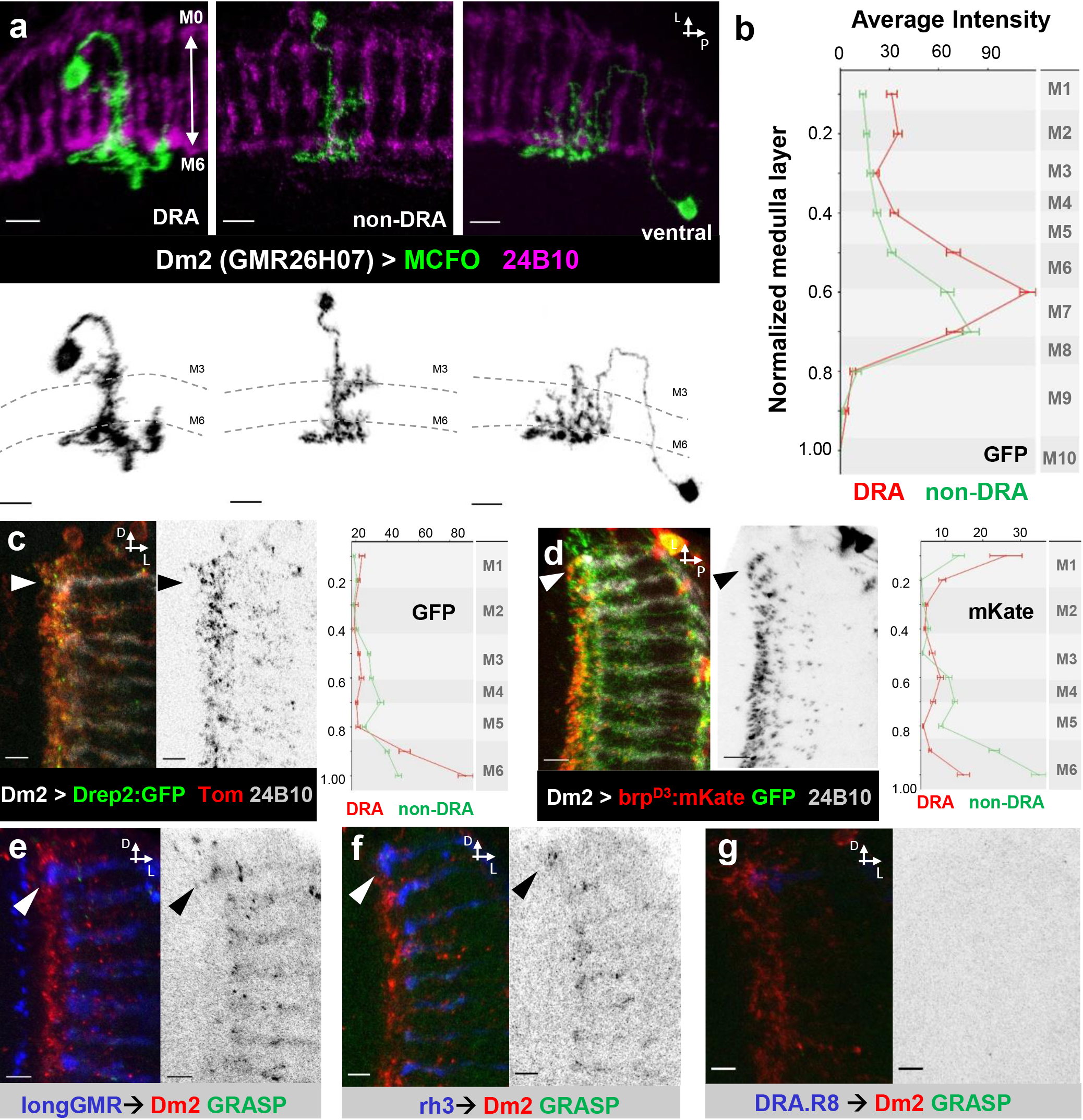
Distal medulla cell type Dm2 displays DRA-specific morphology. **a** Top: Dorsal view of representative MCFO-induced single cell clones of Dm2 cells (green) in the DRA region (left), in the middle (middle) and located at the ventral rim (right) of the medulla (labeled in purple using 24B10). Bottom: grayscale images of the same clones. **b** Average GFP intensity of Dm2 cells located within DRA columns (red) versus non-DRA columns (green) plotted along the distal-proximal axis of the medulla ranging from layer M1-M10 (for layer annotation see methods). **c** Left: Putative postsynaptic membranes (visualized through expression of Drep2::GFP fusion proteins, in green) of Dm2 cells with corresponding grayscale image (middle). Right: quantitative analysis of the distribution of GFP signals in the DRA region (red) compared to non-DRA columns (green), spanning from layer M1-M6. **d** Left: Putative presynaptic membranes of Dm2 cells (visualized through expression of BRP^D3^::mKate fusion proteins, in red) with corresponding Greyscale image (middle). Right: quantitative analysis mKate distribution in the DRA region (red) compared to non-DRA columns (green) spanning from layer M1-M6. **e** Left: Activity-dependent GRASP experiment identifying putative synaptic contacts (in green) between all Photoreceptors (longGMR, in blue) and Dm2 cells (in red). Right: Single channel grayscale image of the GRASP signal. **f** Left: Activity-dependent GRASP identifying putative synaptic contacts (green) between rh3-expressing Photoreceptors (DRA.R7+R8 and pR8, in blue) and Dm2 cells (red). Right: single channel grayscale image of the GRASP signal. **g** Left: Activity-dependent GRASP identifying putative synaptic contacts (green) between DRA.R8 Photoreceptors (blue) and Dm2 cells (red). Right: single channel grayscale image of the GRASP signal. Scale bars: 7μm in (a), 5 μm in (c-g).

We next visualized single cell clones of Dm9 cells in the DRA, since this cell is believed to be a major synaptic target of both R7 and R8, while providing synapses back onto these photoreceptors (Heath, Christenson et al. 2019). To our surprise, Dm9 morphology differed only slightly between DRA and non-DRA columns, reaching into slightly deeper layers in the former (Fig. 3a,b). Importantly, a given clone located at the dorsal rim shared photoreceptor contacts with both DRA and non-DRA columns, pointing towards this cell type not being modality-specific (Fig. 3c). The distribution of putative post-synaptic membranes of Dm9 cells in the DRA showed higher intensity in layers M2-M3, slightly more distally when compared to non-DRA columns, where signal was elevated in layer M4(DRep2:GFP; Fig. 3d). Presynaptic signal in DRA columns remained elevated from M3 throughout M6 (where it weakened), whereas the signal in non-DRA columns formed a distinct peak in M3 which was separated from a weaker peak in M6 (Fig 3e). Finally, using activity-dependent GRASP we observed reconstitution of GFP signals, both when Dm9 cells were pre- or postsynaptic to photoreceptors, in DRA as well as in non-DRA medulla columns (Fig 3f,g). It must be noted that all other Dm cell types analyzed showed no obvious differences between DRA and non-DRA columns (for example Dm4, see Supplemental Fig. S2).

**Fig. 3:**
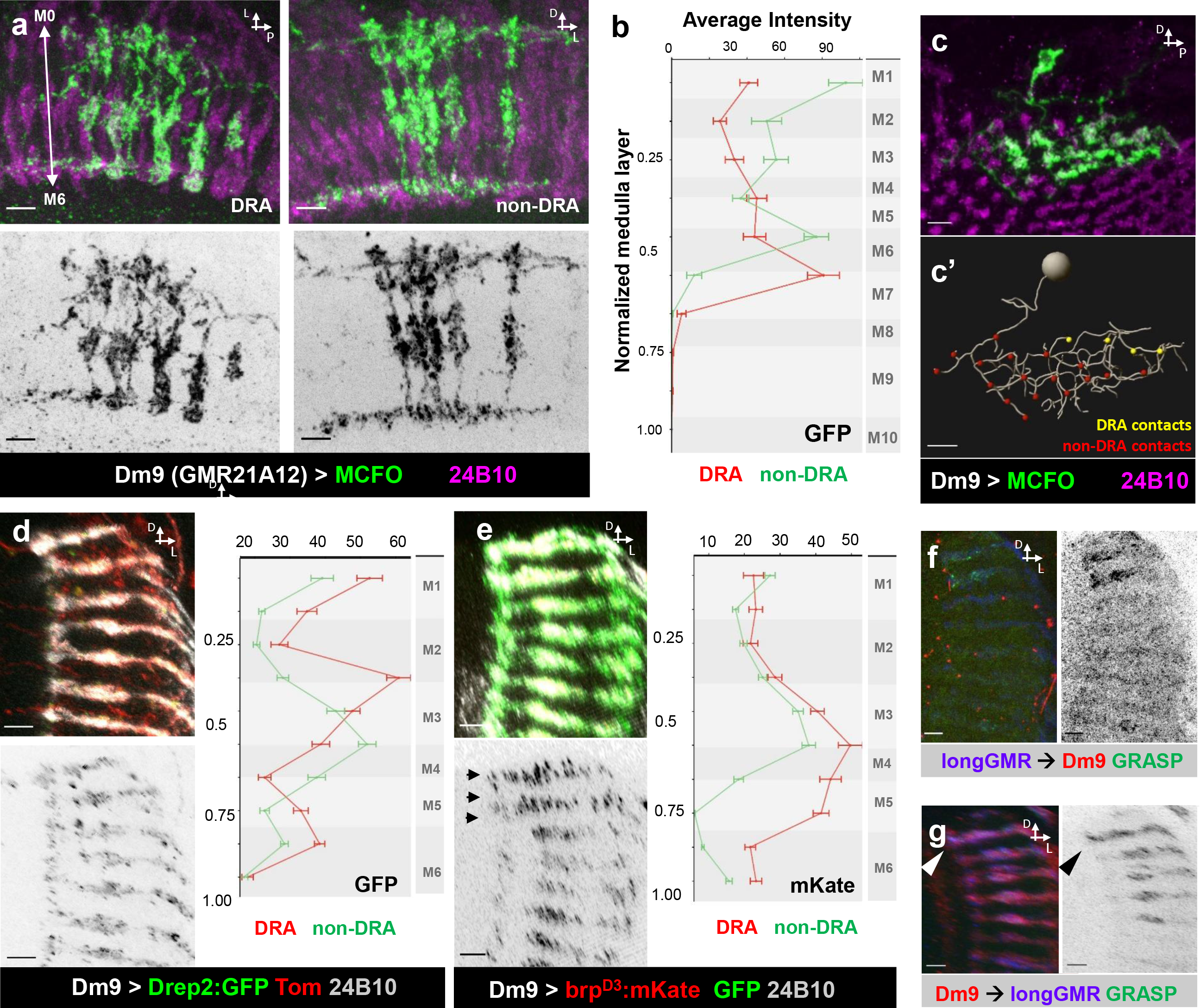
Distal medulla cell type Dm9 displays DRA-specific presynaptic specializations. **a** Top: Dorsal view of representative MCFO-induced single cell clones of Dm9 cells (green) in the DRA region (left), and located outside (right); medulla labeled in purple using −24B10. Bottom: grayscale images of the same clones. **b** Average GFP intensity of Dm9 clones located within DRA columns (red) versus non-DRA columns (green) plotted along the distal-proximal axis of the medulla ranging from layer M1-M10 (for layer annotation see methods). **c** Lateral view of a Dm9 single cell clone (green) located at the dorsal rim of the medulla. **c’** Skeleton of the above cell with DRA photoreceptor contacts in yellow and non-DRA contacts shown in red. **d** TOP: Putative postsynaptic membranes (visualized through expression of Drep2::GFP fusion proteins, in green) of Dm9 cells (labeled with UAS-Tomato, in red) with corresponding grayscale image of the GFP signal (bottom). All photoreceptors are labeled using Anti-24B10 (Chaoptin, in grey). Right: quantitative analysis of the distribution of GFP signals in the DRA region (red) compared to non-DRA columns (green), spanning from layer M1-M6. **e** Top: Putative presynaptic membranes (visualized through expression of BRP^D3^::mKate fusion proteins, in red) of Dm9 cells (labeled with UAS-mCD8:GFP, in green) with corresponding Greyscale image of the mKate signal (bottom). All photoreceptors are labeled using Anti-24B10 (Chaoptin, in grey). Right: quantitative analysis mKate distribution in the DRA region (red) compared to non-DRA columns (green) spanning from layer M1-M6. **f** Left: Activity-dependent GRASP experiment identifying putative synaptic contacts (in green) between all Photoreceptors (presynaptic, labeled with longGMR in blue) and Dm9 cells (postsynaptic, in red). Right: Single channel grayscale image of the GRASP signal. **g** Left: Activity-dependent GRASP experiment identifying putative synaptic contacts (in green) between Dm9 cells (presynaptic, in red) and all photoreceptors (postsynaptic, labeled with longGMR in blue). Right: Single channel grayscale image of the GRASP signal. Scale bars: 5μm.

### Morphology and connectivity of different transmedulla (Tm) cell types in the DRA

To confirm that, like in other insects, polarized light information is transmitted to the lobula complex, we next visualized single cell clones of different transmedulla (Tm) cell types, starting with Tm5c, one out of three Tm5 subtypes (Meinertzhagen, Takemura et al. 2009), which is known to be the main synaptic target of non-DRA R8 cells (Gao, Takemura et al. 2008, Karuppudurai, Lin et al. 2014). No significant morphological differences were seen between Tm5c cells in the DRA, or elsewhere in the medulla (Fig 4a,b). However, both the distribution of post- and presynaptic signals (DRep2:GFP and brp^D3^:mKate, respectively) were altered in DRA columns, spreading towards deeper medulla layers (Fig. 4c). Interestingly, activity-dependent GRASP between DRA R7+R8 and Tm5c (Fig 4d) versus only DRA.R8 and Tm5c (Fig 4e) clearly identified GFP reconstitution between Tm5c as a potential DRA.R7 target (with signal detected between rh3-expressing photoreceptors and Tm5c cells, but no signal detectable between DRA.R8 cells and Tm5c). Similarly, our experiments performed for another known R8 target, the transmedulla cell type Tm20 also revealed no morphological differences between DRA and non-DRA columns, yet revealed no synaptic connections between R7 or R8 in the DRA (Supplemental Fig. S3). However, this could also be due to weak and variable expression observed for this particular Gal4 driver. To our surprise, expression of a driver specific to the two remaining subtypes of Tm5 cells (Tm5a,b) (Fig. 5a,b) was specifically absent from DRA columns (Fig 5c,d), raising the possibility that this cell type is not present in DRA columns.

**Fig. 4:**
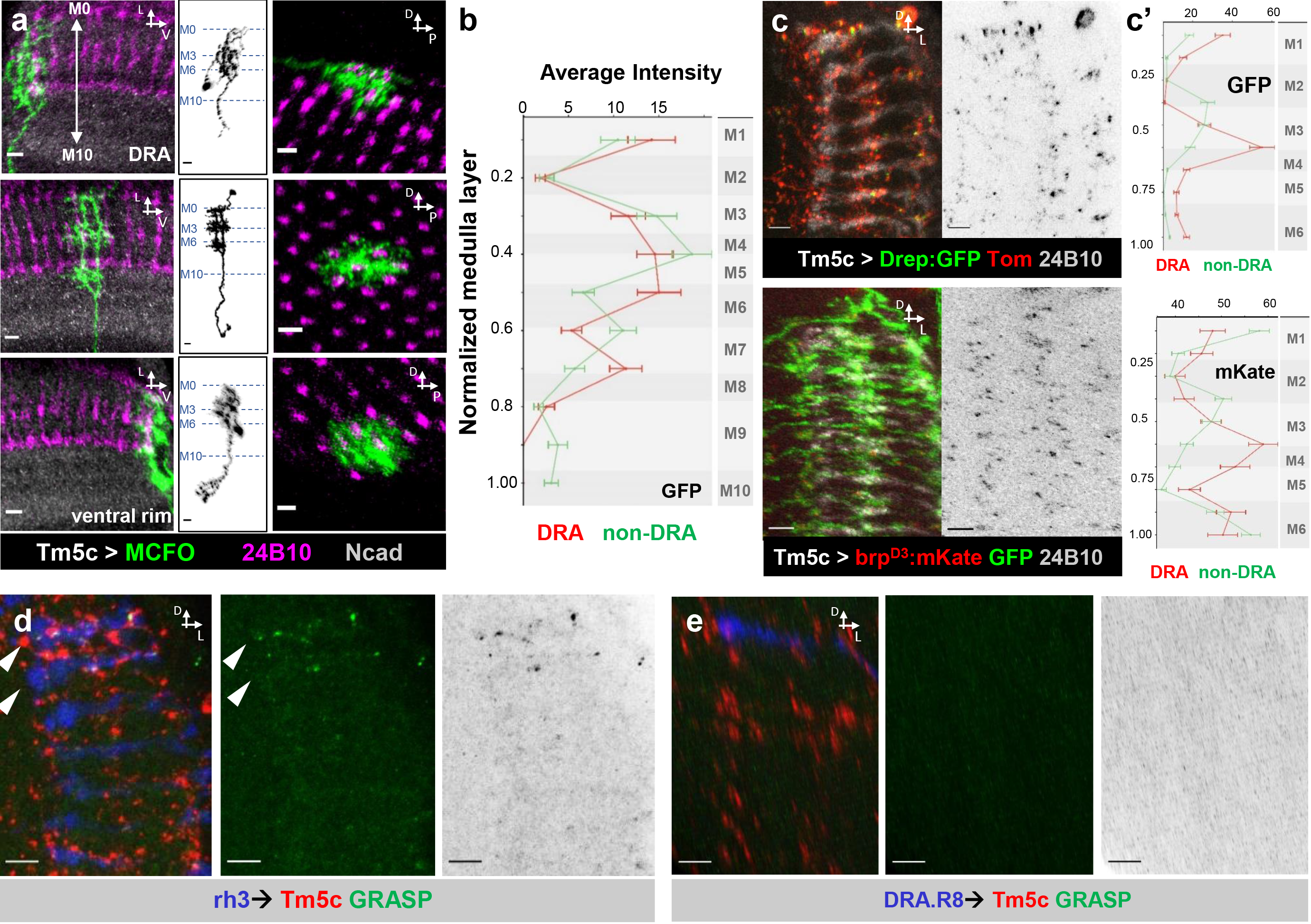
Morphology of the transmedulla cell type Tm5c in the DRA and beyond. **a** Representative MCFO-induced single cell clones of Tm5c cells (green) in the DRA region (first row), in the middle of the medulla (middle row) and located at the ventral rim (bottom row). Anterior view of individual Tm5c cells within the medulla (labeled in purple using 24B10 and anti-Ncad in grey; first column), grayscale image of the same clones (second column) and their lateral view (third column). **b** Average GFP intensity of Tm5c clones located within DRA columns (red) versus non-DRA columns (green) plotted along the distal-proximal axis of the medulla ranging from layer M1-M10 (for layer annotation see methods). **c** Top row: Putative postsynaptic membranes (visualized via expression of Drep2::GFP fusion proteins, in green) of Tm5c cells (visualized using UAS-Tomato, in red), with corresponding Greyscale images of the GFP signal to the right. Bottom row: presynaptic membranes (visualized by the expression of BRP^D3^::mKate fusion proteins, in red) of Tm5c cells (visualized using UAS-mCD8:GFP, in green) with corresponding Greyscale images of the mKate signal to the right. In both cases, all photoreceptors were labeled using Anti-24B10 (Chaoptin), in grey. c’: quantitative analysis GFP (top) and mKate (bottom) distribution in the DRA region (red) compared to non-DRA columns (green) spanning from layer M1-M6. **d** Left: Activity-dependent GRASP experiment identifying putative synaptic contacts (in green) between rh3-expressing Photoreceptors (DRA.R7+R8 and pR8, in blue) and Tm5c cells (in red). Middle: Single channel image of the GRASP signal and corresponding grayscale image (right). **e** Activity-dependent GRASP identifying putative synaptic contacts (green) between DRA.R8 Photoreceptors (blue) and Tm5c cells (red). Middle: single channel images of the GRASP signal and corresponding grayscale image (right). Scale bars: 5μm.

**Fig. 5:**
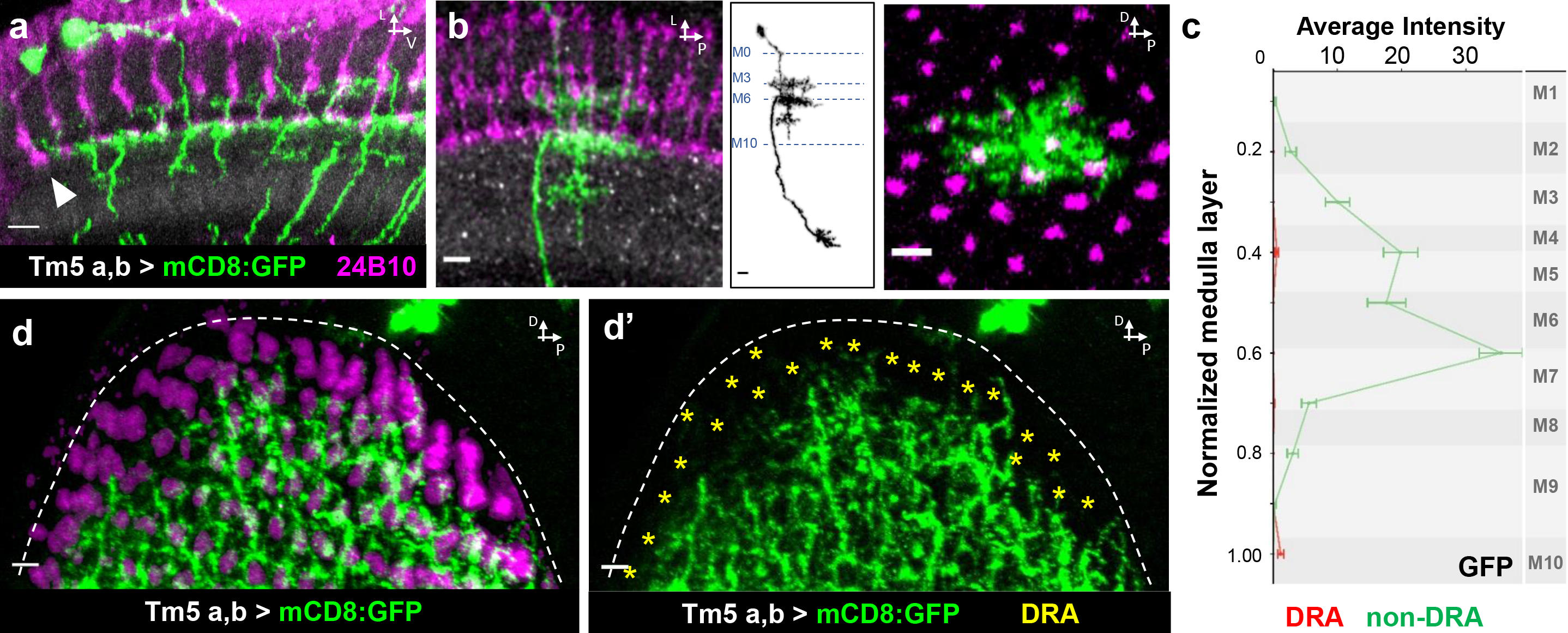
Absence of transmedulla cell types Tm5a,b from the DRA region. **a** Expression of membrane tethered GFP using Tm5a,b-Gal4 reveals an absence of GFP signal in the DRA (arrowhead). **b** Representative MCFO-induced single cell clone of Tm5a,b (green) and single channel grayscale image (middle), as well as lateral view (right). **c** Average Tm5c GFP signal intensity from DRA columns (red) versus non-DRA columns (green) plotted along the distal-proximal axis of the medulla ranging from layer M1-M10 (for layer annotation see methods). **d** Lateral view of medulla expression of Tm5a,b-Gal4 reveals an absence of GFP signal from virtually all DRA columns (yellow asterisks). Scale bars: 5μm.

### Morphology and connectivity of different multicolumnar cell types in the DRA

The medulla neuropil alone contains at least 80 cell types, many of which span multiple columns (Fischbach and Dittrich 1989). We therefore wondered whether some of these cell types would specifically avoid mixing information from DRA and non-DRA columns. Indeed, we identified one Gal4 line expressed in cells stratifying below layer M6, spanning the entire medulla in close vicinity to R7 terminals (Fig 6a). Interestingly, photoreceptor contacts were specifically avoided in DRA columns, where GFP signal was clearly separated from photoreceptor terminals, whereas signals overlapped in the non-DRA region (Fig. 6a,b). Multicolor single cell clones identified the labeled cells as medulla tangential cells similar to Mt11 (Fischbach and Dittrich 1989) manifesting receptive fields of varying sizes, increasing in size towards ventrally (Fig. 6c) and processes passing by the lobula and terminating in the ventrolateral protocerebrum (VLP). More careful analysis revealed that two different cell types were labeled by this driver line, which we termed ‘single layer cells’ and ‘double layer cells’, due to their dendritic morphology in the medulla (Fig. 6d). Interestingly, only single layer cells (Mt11-like cells) specifically avoided contacts with photoreceptor cells from DRA columns (Fig. 6e,f).

**Fig. 6:**
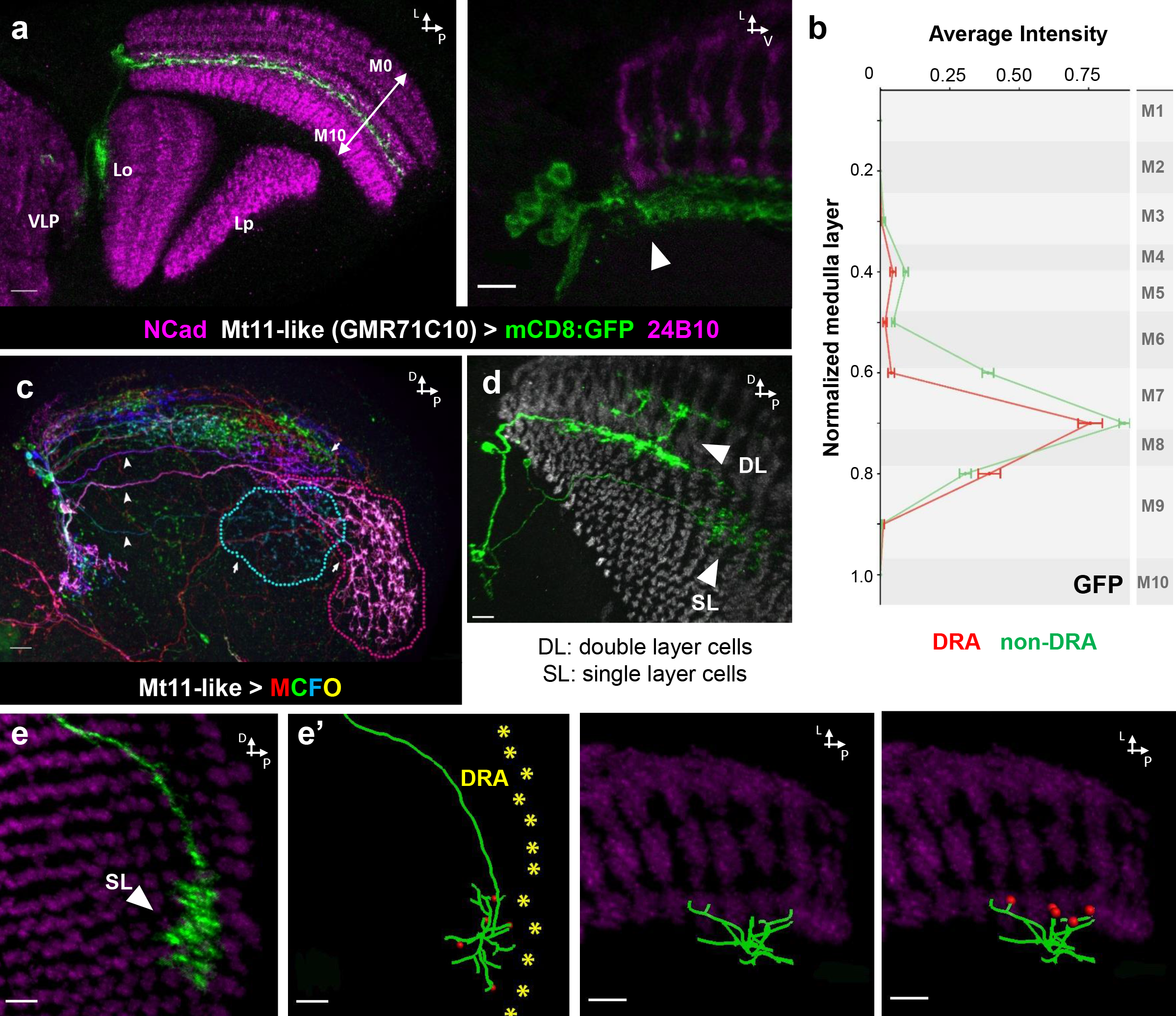
Mt11-like medulla tangential cells specifically avoid the DRA region. **a** Left: Expression of membrane tethered GFP using GMR71C10-Gal4 expressed in Mt11-like medulla tangential cells (green) with processes bypassing the lobula (Lo) and terminating in the ventrolateral protocerebrum (VLP). Lp = lobula plate. Right: Magnification of the dorsal rim reveals absence of GFP signal in the DRA, below layer M6 (arrowhead). **b** Average Mt11-like GFP signal intensity from DRA columns (red) versus non-DRA columns (green) plotted along the distal-proximal axis of the medulla ranging from layer M1-M10 (for layer annotation see methods). **c** Multi-color labeling of Mt11-like cells reveals individual cell morphologies tiling the medulla, with processes (arrow heads) extending to dendritic fields of varying sizes, with largest receptive fields (arrow) located ventrally (dashed lines). **d** Two single cells clones obtained using GMR71C10-Gal4 indicative of distinct cell types labeled: single layer cells (SL) and double layer cells (DL). **e** Left: Lateral view of an Mt11-like single cell clone (green) avoiding contacts with the dorsal rim of the medulla. Right: Skeleton of the same cell with DRA columns labeled with yellow asterisks, photoreceptor contacts labeled with red balls (e’). **f** Left: Dorsal view of the skeleton of the above Mt11-like single cell clone (green) avoiding photoreceptor contacts with the dorsal rim of the medulla. Right: contacts with non-DRA photoreceptors labeled with red balls (f’). Scale bars: 10μm in (a),(c),(d) and 5μm in (b) (e).

We then turned to even larger neuromodulatory neurons, spanning many medulla columns, up to the entire neuropil. Interestingly, octopaminergic neurons labeled by Tdc2-Gal4 (Cole, Carney et al. 2005) (Fig. 7a) also specifically avoided layer M6 specifically in the DRA region (Fig. 7b,c). This effect seemed to be rather specific for octopamine since other neuromodulatory cell types (Friggi-Grelin, Coulom et al. 2003) did not show the same phenotype (Supplementary Fig. S4).

**Fig. 7:**
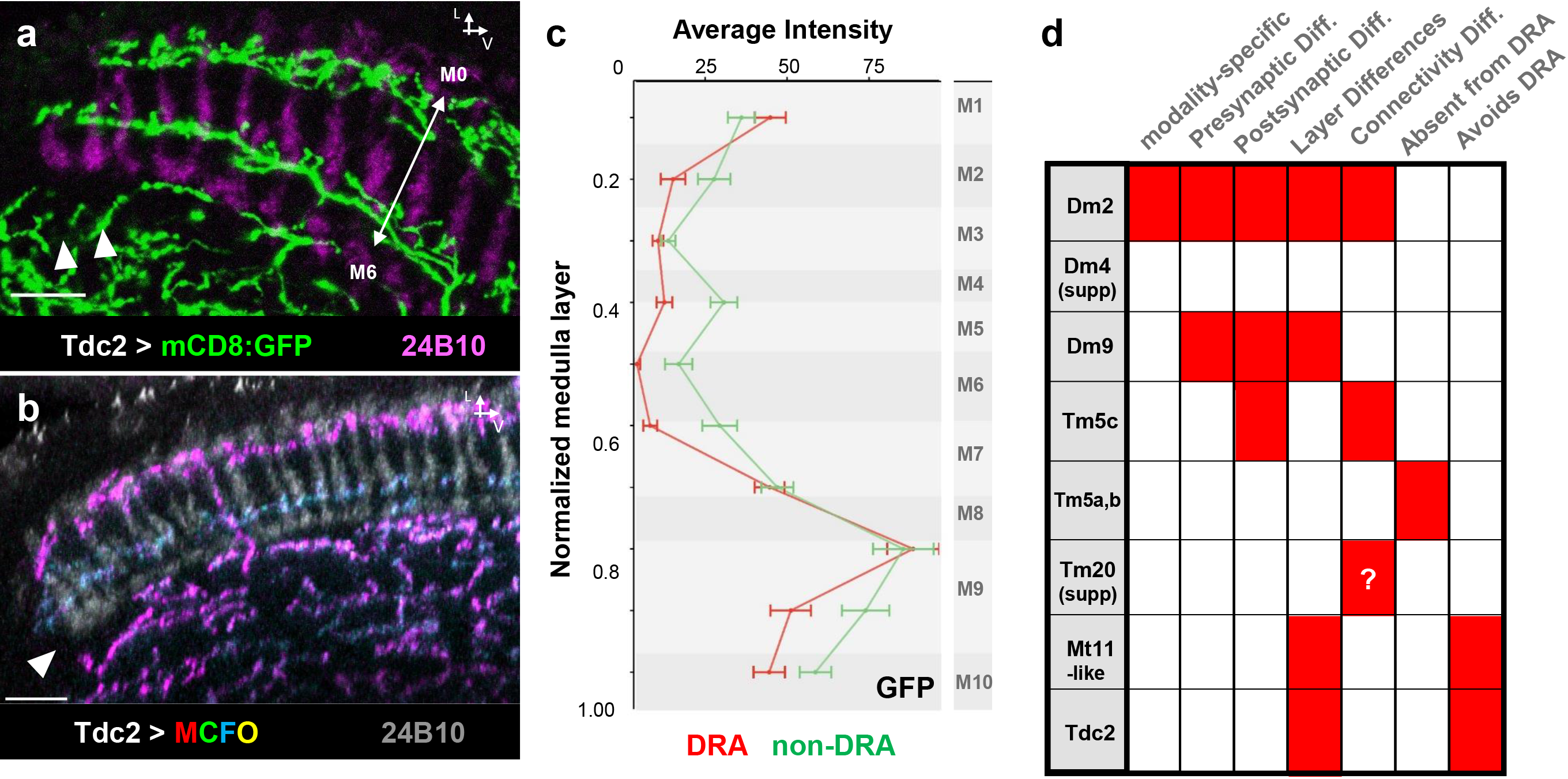
Octopaminergic modulatory neurons avoid layer M6 in the DRA region. **a** Expression of membrane tethered GFP using Tdc2-Gal4 for labeling octopaminergic neurons in the medulla reveals a specific absence of GFP signal in the layer M6 of the DRA (arrowheads). **b** Multi-color labeling of Tdc2-expressing single cell clones reveals at least two populations of cells, one of which avoids layer M6 in the DRA (purple), whereas the other covers the entire layer M3/4, both in the DRA as well as beyond (blue). **c** Average Tdc2-expressing GFP signal intensity from DRA columns (red) versus non-DRA columns (green) plotted along the distal-proximal axis of the medulla ranging from layer M1-M10 (for layer annotation see methods). **d** Summary of DRA-specific features of all cell types analyzed in this study. Scale bars: 10μm.

## Discussion

In the insect optic lobes, repetitive columnar microcircuits post-synaptic to photoreceptor cells process visual information. The goal of this study was to characterize differences in morphology and synaptic connectivity (by comparison of activity GRASP signals between DRA and non-DRA columns) of neuronal elements located in medulla columns downstream of functionally specialized DRA ommatidia that detect the celestial polarization pattern in order to inform orientation and navigation responses. Only one of the cell types analyzed here (Dm2) appeared to be modality-specific in a ways that we had previously demonstrated for Dm-DRA1 and Dm-DRA2 cells (Sancer, Kind et al. 2019), i.e. specifically contacting only polarizationsensitive inputs, while avoiding contacts with non-DRA inputs (summarized in Fig. 7d). Surprisingly, our data points towards Dm2 being a potential synaptic target of DRA.R7 cells (as determined via activity GRASP), whereas Dm2 has previously been shown to be a target of non-DRA R8 photoreceptors (Takemura, Bharioke et al. 2013). From the DRA-specific differences in dendritic morphology, synaptic distribution, and putative connectivity presented here, Dm2 cells in the DRA might form a discrete cell type related to, but different from non-DRA Dm2 cells (in analogy to Dm-DRA cells and Dm8 cells). For now, this cannot be conclusively proven, due to the lack of transcription factors specifically labeling these cells. Interestingly, we observed the same switch in putative R7 vs R8 specificity for Tm5c cells. The reason for such a potential switch of synaptic partners remains unknown. However, it should be pointed out that in adult fly DRA ommatidia, both inner photoreceptors assume an R7-like fate, expressing an R7 Rhodopsin (Rh3) (Fortini and Rubin 1990), DRA.R8 losing expression of the R8-specific transcription factor Senseless (Wernet, Labhart et al. 2003, Wernet and Desplan 2014), both cells sending axons to layer M6 (Strausfeld and Wunderer 1985, Fischbach and Dittrich 1989, Chin, Lin et al. 2014, Sancer, Kind et al. 2019), and assuming an R7-like distribution of presynaptic sites (Sancer, Kind et al. 2019). Due to this similarity, a functional distinction between R7 and R8 becomes increasingly difficult in the DRA and it therefore seems plausible that they share similar post-synaptic elements there. Surprisingly, we found that Dm9 cells not to be modality-specific, meaning that they either mix polarization and colour information, or maybe these cells process visual information that is not related to these qualities, like overall intensity between medulla columns.

Most cell types analyzed here show DRA-specific changes in the distribution of post- or presynaptic membranes. In most cases, this reflects the fact that only in the DRA, R8 cells target to the deeper layer M6, thereby pushing R8-specific signals to the deeper levels (Dm2, Dm9, Tm5c). Beyond this, several cell types show more specific differences, like an absence of signal in layer M3 (Dm2), or a more uniform signal without any gaps usually observed in non-DRA columns, extending continuously from layers M3 to M6 (Dm9). Overall, these DRA-specific changes in pre- and postsynaptic density distribution are a clear indication of modality-specific differences in synaptic connectivity or synaptic strength within DRA ommatidia, not only between photoreceptors and the cell types studies, but potentially involving additional cell types many of whose cell type identity may currently remain unknown. Our recent study showed that Dm-DRA1 and Dm-DRA2 cells integrate from ~10 neighboring DRA.R7 cells (or DRA.R8 cells, respectively), thereby revealing an important difference for how R7 and R8 signals are being compared in DRA versus non-DRA ommatidia. The changes in pre- and postsynaptic signals described here now provide an additional example for how DRA-specific differences in synaptic circuitry may shape local computations (polarization versus color).

Of all the transmedulla cell types tested here (Tm5a,b, Tm5c, Tm20), only Tm5c is likely to be synaptically connected to polarization-sensitive photoreceptors (presumably DRA.R7, as determined indirectly via activity-dependent GRASP), revealing that polarized light information might be represented in the lobula neuropil. This is in agreement with studies from locusts (Homberg 2015), as well as with our previous trans-synaptic tracing study (Sancer, Kind et al. 2019), which independently revealed Tm cells post-synaptic to DRA.R8 cells. As the same experiment also revealed potentially direct connections between medulla and Anterior Optic Tubercle (AOTU), it appears that polarized light information is processed via several – possibly interconnected – pathways. It remains unknown how many different Tm cell types connect the DRA to the lobula, and the absence of Tm5a,b cells from DRA columns suggests that this pathway might require a rather specific complement of neuronal elements. In agreement with this, we also observed medulla tangential cells (Mt11-like) specifically avoiding DRA columns, while tiling across the entire rest of the medulla. Although the functional role of these cells remains unknown it seems clear their target area in the ventrolateral protocerebrum (VLP) appears to collect information from the entire medulla, yet specifically lacking polarized light inputs from the DRA, indicating that different visual qualities in represented separate areas in the central brain.

Finally, the avoidance of specific DRA medulla layers by large octopaminergic neurons suggests that neuromodulation may affect differently those visual circuits computing different visual modalities. Opposing effects of both octopamine and dopamine on visual behavior have previously been demonstrated (Gorostiza, Colomb et al. 2016), and it remains to be seen how these neuromodulators affect navigation behaviors by modulating the underlying circuitry.

## Acknowledgements

The authors would like to thank Stephan Sigrist and Robin Hiesinger for sharing fly stocks and reagents, as well as two anonymous reviewers who made helpful suggestions on the manuscript. This work was supported by the Deutsche Forschungsgemeinschaft (DFG) through grants WE 5761/2-1 and SFB958 (Teilprojekt A23), through AFOSR grant FA9550-19-1-7005, through the Berlin Excellency Cluster NeuroCure, with support from the Fachbereich Biologie, Chemie & Pharmazie of the Freie Universität Berlin, as well as the Division of Neurobiology at Freie Universität Berlin (support of FU Berlin and the National Institute of Health to Robin Hiesinger).

