## Supplemental Data for "Cellular and synaptic adaptations of neural circuits processing skylight polarization in the fly"

**Supplemental Tables:**

- **Supplemental Table1: Abbreviations used in the text**
- **Supplemental Table 2: List of cell types**

**Supplemental Figures:**

- **Supplemental Fig. S1: Dm2 cells are modality-specific**
- **Supplemental Fig. S2: Morphology of Dm4 cells in the DRA and beyond**
- **Supplemental Fig. S3: Morphology of Tm20 cells in the DRA and beyond**
- **Supplemental Fig. S4: Dopaminergic neurons do not respect the DRA boundary**

**Supplemental Table1: Abbreviations used in the text**

| | |
|---|---|
| 24B10 | Monoclonal antibody against Chaoptin (photoreceptor marker) |
| A | Anterior |
| AD | Activation domain (split GAL4) |
| AOTU | Anterior Optic Tubercle |
| B | Blue |
| BDSC | Bloomington Drosophila Stock Center |
| BRP | Bruchpilot (marker of presynaptic active zones), as in brpD3 (short fragment) |
| D | Dorsal |
| DBD | DNA-binding domain (split GAL4) |
| Dm | Distal medulla (designation of cell type), as in Dm2, Dm4, Dm8, Dm9 |
| DRA | Dorsal Rim Area |
| DRA.R7 | R7 photoreceptor cell from DRA ommatidia |
| DRA.R8 | R8 photoreceptor cell from DRA ommatidia |
| DRep2 | Marker of putative postsynaptic membranes |
| eq | Equator of the eye |
| FLP | Flip recombinase ('flippase') |
| G | Green |
| GAL4 | Transcription factor (binary expression system) |
| GFP | Green fluorescent protein |
| GMR | Glass multimeric reporter, as in longGMR |
| GRASP | GFP reconstitution across synaptic partners (technique) |
| L | Lateral |
| La | Lamina (neuropil) |
| LexA | Transcriptional activator recognizing LexAop binding sites |
| Lo | Lobula (neuropil) |
| Lp | Lobula plate (neuropil) |
| M | Medulla layer, as in M3 or M6 |
| mCD8:GFP | Membrane-tagged GFP |
| MCFO | Multi-color Flp out (technique) |
| Me | Medulla (neuropil) |
| mKate | Red-fluorescent protein |
| Mt | Medulla tangential (cell type) |
| NCad | N-Cadherin (neuropil marker) |
| PBS | Phosphate buffered saline |
| R | Photoreceptor, as in R7 or R8 |
| Rh | Rhodopsin, as in rh3 |
| Tdc | tyrosine decarboxylase |
| TH | Tyrosine hydroxylase |
| Tm | Transmedullary (cell type), as in Tm5a,b or Tm5c, Tm20 |
| Tom | Tomato (red fluorescent protein) |
| UAS | Upstream activated sequences (recognized by GAL4) |
| UV | Ultraviolet |
| VLP | Ventrolateral protocerebrum |

**Supplemental Table 2: List of cell types**

| | Cell type | Description | Connectivity | | Reference |
|---|---|---|---|---|---|
| | | | R7 | R8 | |
| Distal medulla cells | Dm2 | Stratifies from M3 to M6. Projects to the next column in M6. They occupy ~2 columns in M6 and one in other layers. | Dm2 is postsynaptic | Very few synapses | ( Fischbach and Dittrich, 1989, Takemura et al. 2013, 2015; Nern et al. 2015) |
| | Dm4 | stratifies from M3 to M5/M6 | No synapses | No synapses | ( Fischbach and Dittrich, 1989, Takemura et al. 2013, 2015; Nern et al. 2015) |
| | Dm9 | Stratifies from M1 to M6. It as a tubular shape, and extends between the M1 and M6 layers | Presynaptic and Postsynaptic | Presynaptic and postsynaptic | ( Takemura et al. 2013, 2015; Nern et al. 2015) |
| Transmedullary cells | Tm5a/b | Tm5a: Single main dendritic branch along photoreceptor terminals and fine processes in M3 and M6. Terminates in the Lobula layer 5. Tm5b: extends 2 or 3 main dendritic branches in M4 and M6, spans ~ 5 columns. Also branches at M8. Terminates in lobula layer 5. | Both Tm5a and Tm5b are postsynaptic | Few synapses | (Karuppudurai et al., 2014, Fischbach and Dittrich, 1989, Lin et al., 2016, Gao et al., 2008) |
| | Tm5c | Single main dendritic branch with fine processes in M3 and M6. Also dendritic arbors at M1 and terminals in both lobula layers 4 and 5 | Few synapses | Postsynaptic | (Fischbach and Dittrich, 1989, Gao et al., 2008, Lin et al., 2016, Karuppudurai et al., 2014) |
| | Tm20 | Tm20 arborizes in medulla layers M1–M4 and M8 and terminates in the layer 5 of the lobula | Few synapses | Postsynaptic | (Morante and Desplan, 2008, Takemura et al., 2013, Fischbach and Dittrich, 1989) (Takemura et al. 2013, 2015; Nern et al. 2015) |
| other | Mt11-like | Medulla tangential cell: Mt11 neurons may cover more than 20 columns and restrict synaptic arborizations below M6 | Unknown | Unknown | (Fischbach and Dittrich, 1989, Chin et al. 2014) |
| other | Tdc | Octopaminergic neurons: Project from central brain to medulla, makes branches in distal and proximal medulla and in lobula. | Unknown | Unknown | Busch et al. 2009 |

#### Supplemental Figure Legends

**Supplemental Fig. S1: Dm2 cells are modality-specific**

**a** MCFO experiment depicting a representative Dm2 cell clone (cyan, arrowhead) located at the dorsal edge of the medulla (DRA as hatched box). Scale bars: 20µm in (a), 10 µm in (a) and (a'').

**Supplemental Fig. S2: Morphology of Dm4 cells in the DRA and beyond**

**a** Top: Representative MCFO-induced single cell clones of Dm4 cells (green) in the DRA region (left) and outside the DRA (right) with grayscale image of the same below. **b** Average GFP intensity of Dm4 cells located within DRA columns (red) versus non-DRA columns (green) plotted along the distal-proximal axis of the medulla ranging from layer M1-M10 (for layer annotation see methods). **c** Putative postsynaptic membranes of Dm4 cells (visualized through expression of Drep2::GFP fusion proteins; left) with corresponding single channel grayscale image (right). **d** Putative presynaptic membranes (visualized by the expression of $BRP^{D3}$::mKate fusion proteins; left) of Dm4 cells with corresponding single channel grayscale image (right). Scale bars: 5 µm

**Supplemental Fig. S3: Morphology of Tm20 cells in the DRA and beyond**

**a** Representative MCFO-induced single cell clones of Tm20 cells (green) in the DRA region (top row) and in the middle (bottom row) of the medulla (labeled in purple using -24B10 and anti-Ncad in grey; first column), grayscale image of the same clones (second column) and their lateral view (third column). **b** Average GFP intensity of Tm20 cells located within DRA columns (red) versus non-DRA columns (green) plotted along the distal-proximal axis of the medulla ranging from layer M1-M10 (for layer annotation see methods). **c** Putative postsynaptic membranes of Tm20 cells (visualized through expression of Drep2::GFP fusion proteins; left) with corresponding single channel grayscale image (right). **d** Putative presynaptic membranes (visualized by the expression of $BRP^{D3}$:mKate fusion proteins; left) of Tm20 cells with corresponding single channel grayscale image (right). **e** Left: Activity-dependent GRASP identifying putative synaptic contacts (green) between all Photoreceptors (expressing LongGMR-Gal4; blue) and Tm20 cells (red). Right: single channel images of the GRASP signal as corresponding grayscale image. Scale bars: 5 µm.

**Supplemental Fig. S4: Dopaminergic neurons do not respect the DRA boundary**

**a** Expression of TH-Gal4 (see materials and methods) driving UAS-mCD8:GFP in dopaminergic neurons in the medulla (DRA and non-DRA). Scale bar: 10 µm.

#### Supplemental Figure S1

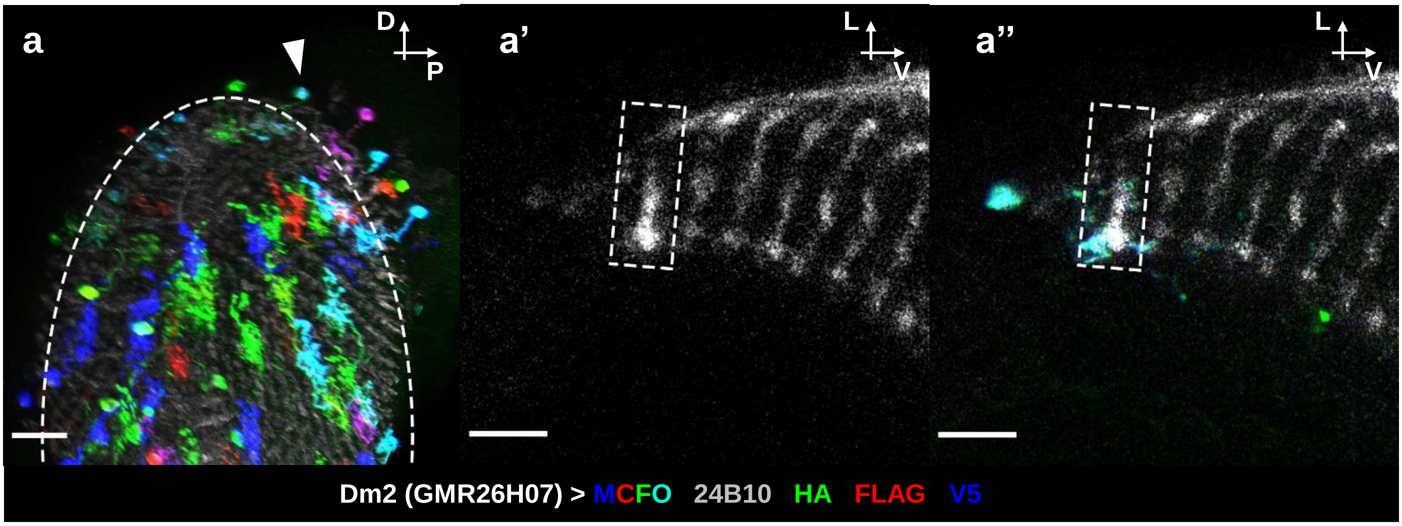

### Supplemental Figure S2

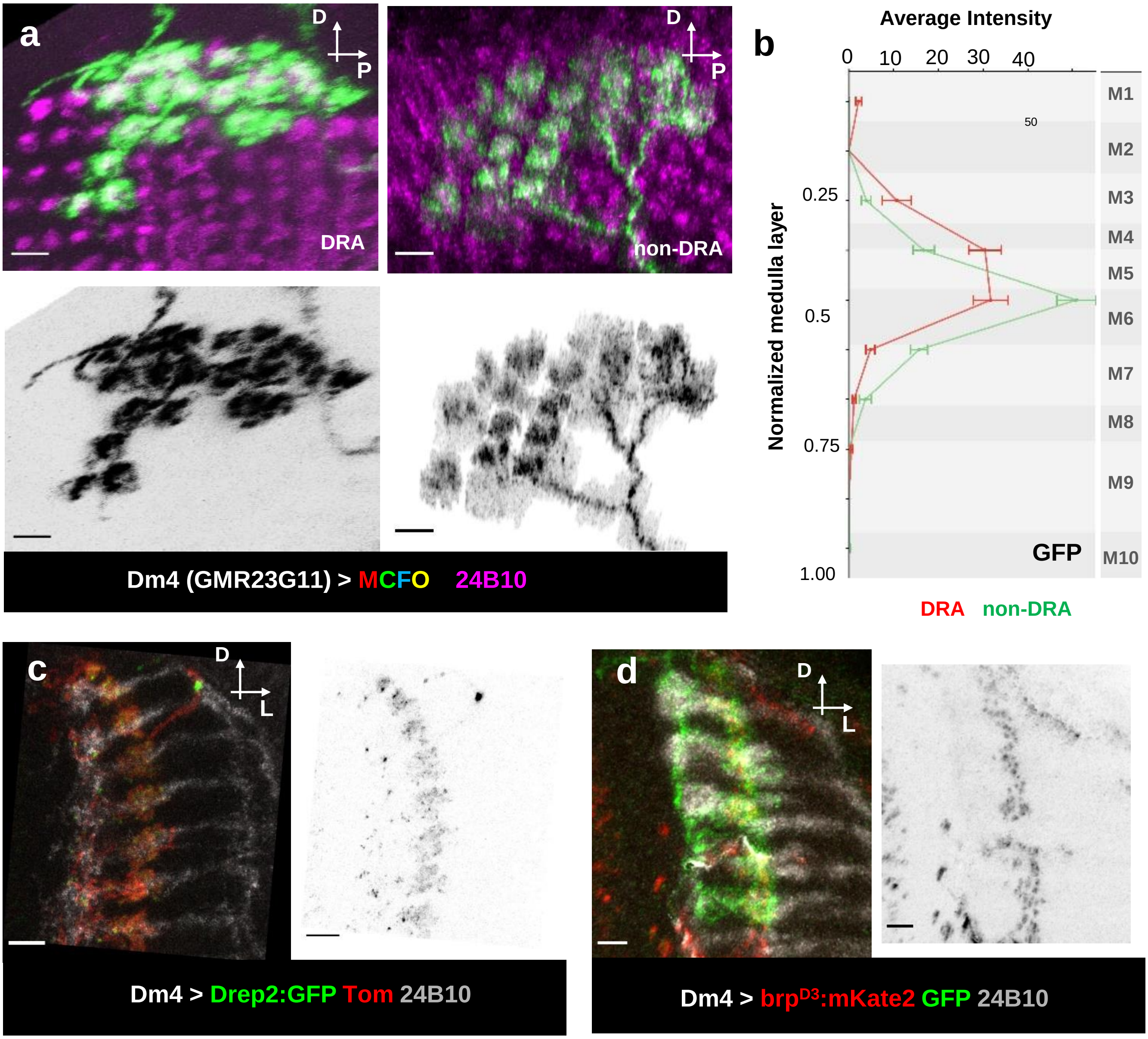

### Supplemental Figure S3

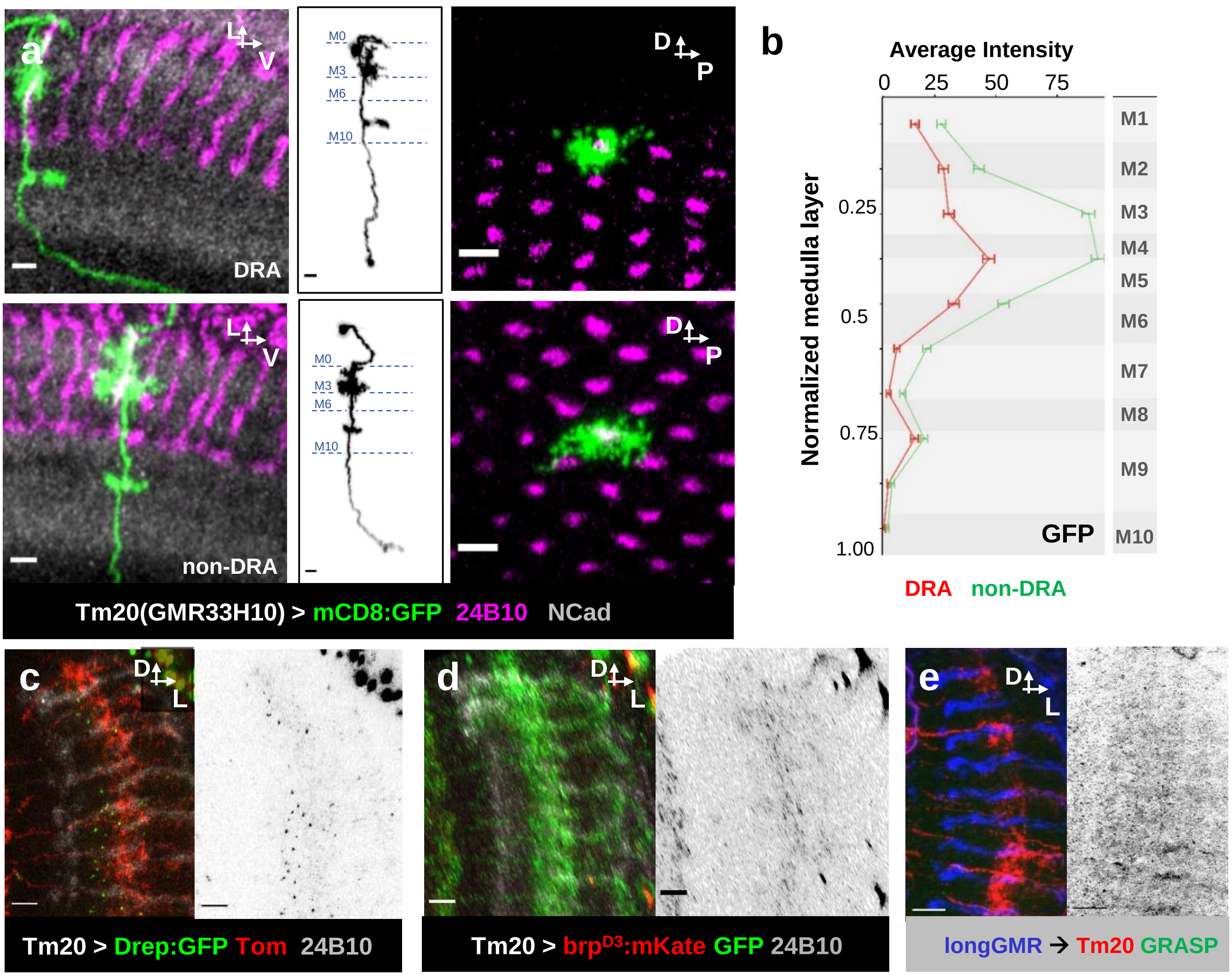

**Supplemental Figure S4**

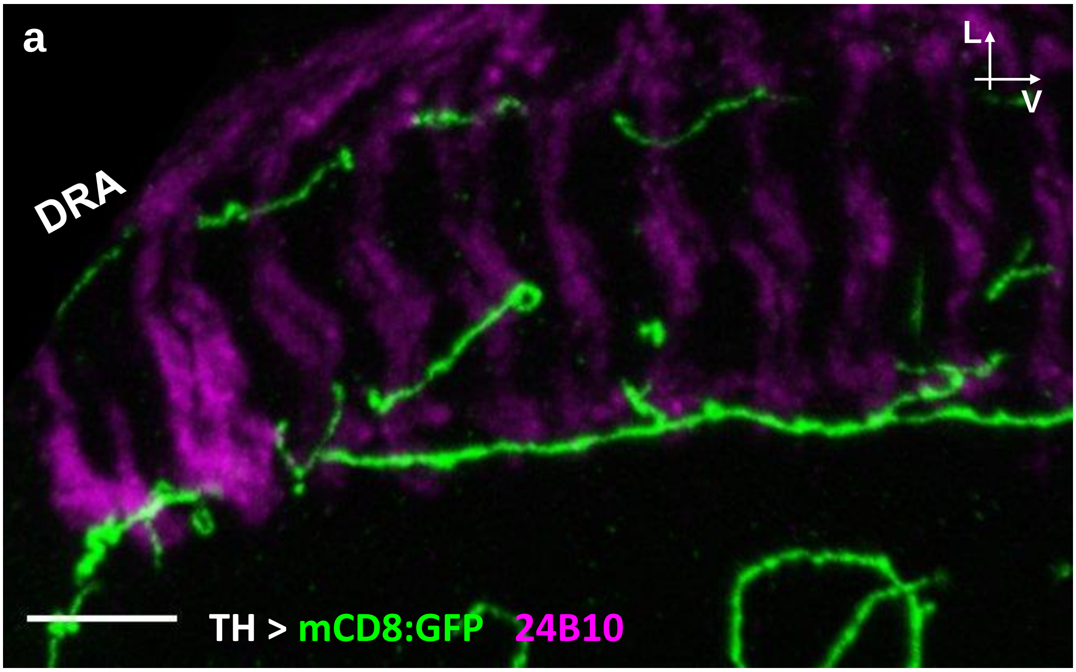
